# Visual working memory recruits two functionally distinct alpha rhythms in posterior cortex

**DOI:** 10.1101/2022.04.15.488484

**Authors:** Julio Rodriguez-Larios, Alma ElShafei, Melanie Wiehe, Saskia Haegens

**Affiliations:** Dept. of Psychiatry, Columbia University, New York, USA, NY 10032; Div. of Systems Neuroscience, New York State Psychiatric Institute, New York, USA, NY 10032; Donders Institute for Brain, Cognition & Behavior, Radboud University, Nijmegen, The Netherlands, 6525 EN

**Author notes:** **Corresponding author:** Julio Rodriguez-Larios, Dept. of Psychiatry, Division of Systems Neuroscience, Columbia University, New York State Psychiatric Institute, 1051 Riverside Drive, Unit 87, New York, NY 10032, USA, **Email:**. **Author contributions** JRL, AE, MW and SH Designed Research. MW Performed Research. JRL, AE and MW Analyzed data. JRL and SH Wrote the paper.

## Abstract

Oscillatory activity in the human brain is dominated by posterior alpha oscillations (8-14 Hz), which have been shown to be functionally relevant in a wide variety of cognitive tasks. Although posterior alpha oscillations are commonly considered a single oscillator anchored at an individual alpha frequency (IAF; ∼10 Hz), previous work suggests that IAF reflects a spatial mixture of different brain rhythms. In this study, we assess whether Independent Component Analysis (ICA) can disentangle functionally distinct posterior alpha rhythms in the context of visual short-term memory retention. Magnetoencephalography (MEG) was recorded in 33 subjects while performing a visual working memory task. Group analysis at sensor level suggested the existence of a single posterior alpha oscillator that increases in power and decreases in frequency during memory retention. Conversely, single-subject analysis of independent components revealed the existence of two dissociable alpha rhythms: one that increases in power during memory retention (Alpha1) and another one that decreases in power (Alpha2). Alpha1 and Alpha2 rhythms were differentially modulated by the presence of visual distractors (Alpha1 increased in power while Alpha2 decreased) and had an opposite relationship with accuracy (positive for Alpha1 and negative for Alpha2). In addition, Alpha1 rhythms showed a lower peak frequency, a narrower peak width, a greater relative peak amplitude and a more central source than Alpha2 rhythms. Together, our results demonstrate that modulations in posterior alpha oscillations during short-term memory retention reflect the dynamics of at least two distinct brain rhythms with different functions and spatiospectral characteristics.

**Significance statement:** Alpha oscillations are the most prominent feature of the human brain’s electrical activity, and consist of rhythmic neuronal activity in posterior parts of the cortex. Alpha is usually considered a single brain rhythm that changes in power and frequency to support cognitive operations. We here show that posterior alpha entails at least two dissociable rhythms with distinct functions and characteristics. These findings could solve previous inconsistencies in the literature regarding the direction of task-related alpha power/frequency modulations and their relation to cognitive performance. In addition, the existence of two distinct posterior alpha rhythms could have important consequences for the design of neurostimulation protocols aimed at modulating alpha oscillations and subsequently cognition.

## Introduction

Working memory entails the storage of information over brief periods of time for its later manipulation (Baddeley, 2010; Repovš & Baddeley, 2006). Traditionally, working memory has been linked to the prefrontal cortex, as neurons in this area show increased spiking when information has to be transiently stored (D’Esposito & Postle, 2015; Fuster & Alexander, 1971). However, more recent research has shown that the brain mechanisms supporting working memory are not limited to the activity of individual neurons in a specific part of the cortex (Miller et al., 2018). Rather, memory traces are distributed in the brain, involving areas beyond the prefrontal cortex (Christophel et al., 2017, 2018). Moreover, the transient maintenance of information involves changes at network level that cannot be well addressed when studying the activity of single neurons (Miller et al., 2018). Neural oscillations, which reflect the summed activity of neural populations (Cohen, 2017), are thought to play an important role in the transient storage of information in the brain (Lundqvist et al., 2016, 2018; Wolinski et al., 2018).

In the human brain, neural oscillations are dominated by alpha rhythms (8–14 Hz) (Bazanova & Vernon, 2014; Hari et al., 1997). Although alpha rhythms are most prominent in posterior areas, they are also found in auditory and sensorimotor cortex (commonly referred to as mu and tau rhythms, respectively) (Bastarrika-Iriarte & Caballero-Gaudes, 2019; Haegens et al., 2011; Lehtelä et al., 1997; Pfurtscheller et al., 2000). Previous work has shown that alpha oscillations desynchronize (decrease in power) in task-relevant areas and synchronize (increase in power) in task-irrelevant areas in a wide variety of cognitive tasks (Haegens et al., 2009; Jensen et al., 2002; Jokisch & Jensen, 2007; Klimesch, 1999). Based on these results, it has been proposed that alpha’s function is to gate information through the brain via functional inhibition (Jensen & Mazaheri, 2010; Klimesch et al., 2007). In line with this idea, recent research in humans demonstrates that the amplitude of alpha oscillations is negatively associated with neural excitability (Chapeton et al., 2019; Haegens et al., 2021; Iemi et al., 2022).

Posterior alpha oscillations are thought to be especially relevant in visual working memory (de Vries et al., 2020). Although several studies have shown significant power modulations of posterior alpha rhythms when visual information has to be transiently stored, the direction of these power modulations is highly inconsistent (Pavlov & Kotchoubey, 2020). Studies reporting increased alpha power during visual memory maintenance argue that this increase is aimed to block visual input by inhibiting (task-irrelevant) occipital and/or parietal areas (Bonnefond & Jensen, 2012; Jensen et al., 2002; Tuladhar et al., 2007). In contrast, studies reporting posterior alpha power decreases during memory retention argue that these occipitoparietal areas are actually task-relevant and need to be disinhibited to support the short-term storage of visual information (De Vries et al., 2018; Erickson et al., 2019; van Ede et al., 2017).

A possible explanation for the inconsistent findings regarding posterior alpha modulations during visual working memory, is the existence of distinct posterior alpha rhythms. Klimesch et al. proposed a division of the alpha band in an upper frequency (∼10-12 Hz) and a lower frequency subcomponent (∼8-10 Hz), based on their differential power modulations during memory tasks (Klimesch, 1999; Klimesch et al., 1993). However, since alpha rhythms were not spatially separated in these studies, we cannot know whether power modulations in the two alpha sub-bands reflect the activity of two different oscillators or a change in frequency of a single oscillator (Haegens et al., 2014; Mierau et al., 2017). The notion of multiple alpha rhythms in posterior areas has been further supported by studies that used source localization techniques (Barzegaran et al., 2017; Benwell et al., 2019; Gulbinaite et al., 2017; Sokoliuk et al., 2019). Benwell et al. (2019) have recently shown that upper and lower alpha rhythms can be spatially disentangled using Independent Component Analysis (ICA), a blind source separation method that allows to identify maximally independent sources of brain activity in M/EEG (Delorme et al., 2012). Nonetheless, the dynamics of these two putative alpha components have not been studied in the context of working memory.

Here, we used ICA to examine whether alpha power modulations in posterior regions during short-term memory retention reflect the activity of one or several brain rhythms. We acquired MEG during a visual working-memory task in which participants (N=33) had to briefly remember one out of four spatial directions, while task difficulty was modulated by introducing visual distractors. We first assessed alpha dynamics at sensor level by comparing alpha power and frequency between fixation and memory retention. Then, we performed the same comparison focusing on occipitoparietal independent components that showed an alpha peak in their spectrum. Interestingly, while results at sensor level suggested a change in power and frequency of a single posterior alpha rhythm during memory retention, our ICA results demonstrated that posterior alpha dynamics reflect the activity of at least two alpha rhythms with distinct functions and spatiospectral characteristics. We believe that these results have important implications not only for the analysis and interpretation of alpha oscillations in M/EEG but also for their potential modulation through different neurostimulation techniques.

## Materials and Methods

### Participants

35 healthy right-handed adult participants (mean age 25.2 years, range 20 to 33; 17 female, 18 male) took part in the experiment. Participants reported normal vision and no history of neurological or mental illnesses. Prior to the experiment, participants were informed about the MEG system as well as safety regulations and signed an informed consent form. The study falls under the general ethics approval (CMO 2014/288 “Imaging Human Cognition”) in accordance with the Declaration of Helsinki. After participation, the participants received a monetary reward. Two participants were excluded due to technical problems during data acquisition.

### Experimental design

MEG was recorded while participants performed a visual working memory task (Figure 1A). This task was designed to emulate real-life situations in which participants would navigate in a city using directions that they had to keep in mind for a short period of time. The goal of the task was to remember one out of four directions. Participants were first presented with a visual direction cue (0.25 s) pointing to the upper left, upper right, bottom left, or bottom right. After a delay period (3 s), a second cue was presented (0.25 s). The second cue was either a ‘stay’ cue, meaning that the correct answer was the direction indicated by the first cue, or a ‘switch’ cue, indicating that the correct answer was the direction opposite to the first cue. After the second cue, a response mapping diagram was shown, indicating which button corresponded to which direction. The correspondence between each button and direction was randomized between blocks (a total of eight different response mapping diagrams were shown). Participants were instructed to answer as quickly and as accurately as possible, with the right hand via a button press. In 50% of the trials, the delay period contained four distractors (randomly drawn direction cues) that were presented at a 0.33 s inter-stimulus interval. Participants were instructed to ignore the distractors and to keep only the first cue in mind. Participants performed eight blocks of 48 trials (∼5 min each), with short breaks between blocks. Participants were seated upright in the MEG helmet and were instructed to keep their head position as stable as possible for the duration of the experiment. Prior to the MEG recording, participants performed a training block (16 trials) to make sure that they understood the task correctly. The experimental stimuli were programmed and presented with the software Presentation (Version 18.0, Neurobehavioural Systems, Inc., Berkeley, CA, www.neurobs.com).

**Figure 1.**
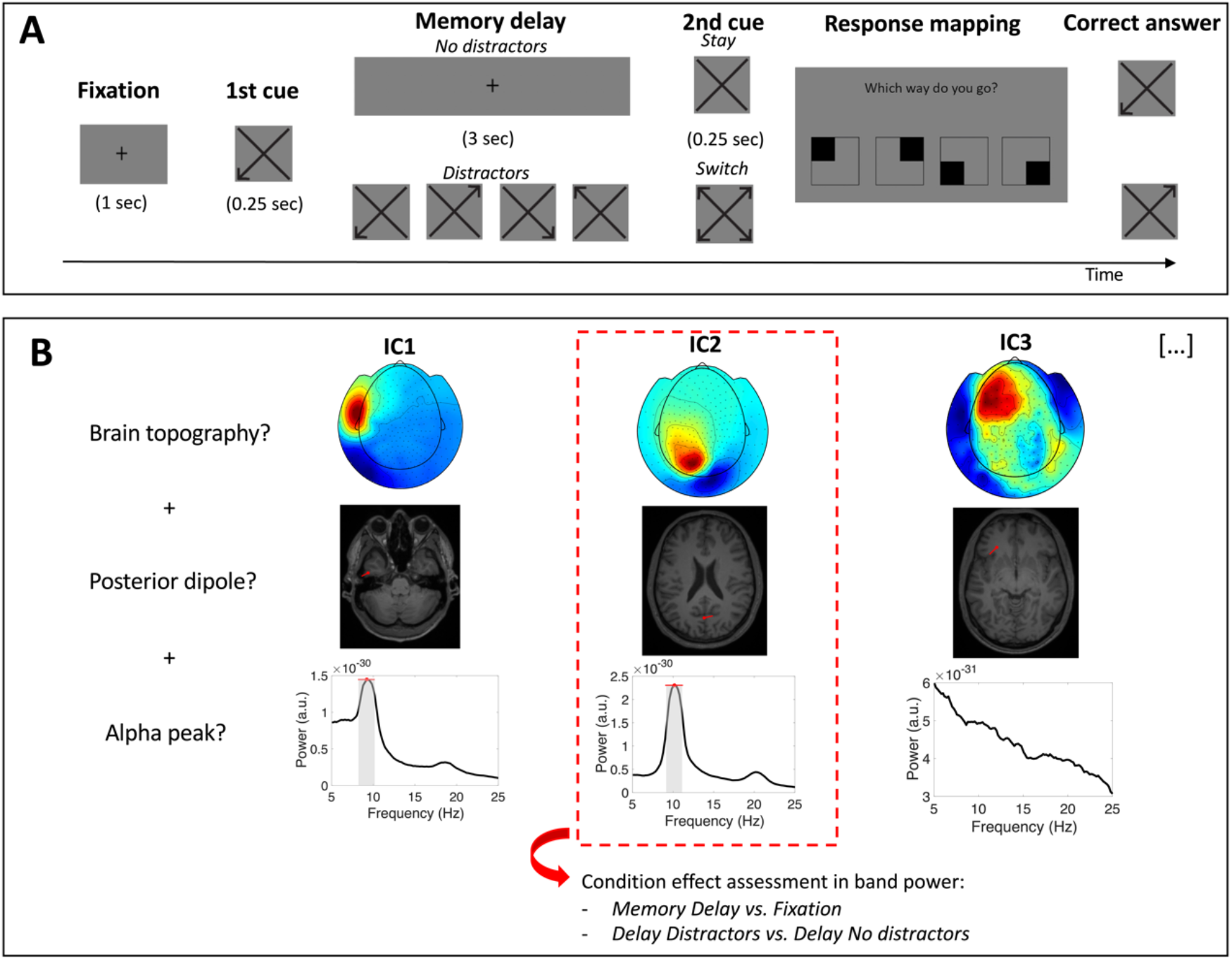
Task and main analytical approach. A) Task schematic. Participants were asked to remember a visual direction cue (i.e., upper left, upper right, bottom left, or bottom right) for 3 s (memory delay). In 50% of the trials, the delay period contained four visual distractors. Based on the content of a second cue (i.e., ‘stay’ or ‘switch’), participants had to report the direction of the first cue or its opposite. B) ICA-based selection in an exemplary subject. A selection of independent components was performed based on their topography (top panel), estimated source (middle), and spectrum (bottom). Only independent components with a brain topography (IClabel algorithm classification > 0.80), an estimated dipole in occipital or parietal cortex and a peak in the alpha range were selected for analysis (in this example, the component marked in the red rectangle). The frequency band of each independent component was defined based on the average spectrum (peak frequency and width; grey area). Condition-related modulations in band power (i.e., delay vs. fixation, distractor vs. no distractor) were assessed through single-trial analysis of each independent component.

### Data Acquisition

MEG data was recorded with a 275-channel CTF MEG system with axial gradiometers at a sampling rate of 1200 Hz (CTF MEG systems, VSM MedTech Ltd.). Six channels were disabled due to permanent malfunction. In order to monitor the head position of the participants and to allow for adjustments to the original position in between blocks, the real-time representation of the participant’s head position was monitored using three head localization coils at the right and left ear canals as well as the nasion (Stolk et al., 2013). These points were further used as offline anatomical landmarks to align the MEG data with structural images of the participant’s brain for source reconstruction. Further, movement of the left eye was tracked during the experiment using an Eyelink eyetracker (SR Research Ltd.). After the experiment, the participant’s head shape was digitized using a Polhemus 3D tracking device (Polhemus, Colchester, Vermont, United States). In a separate session, an anatomical MRI scan of the participant’s brain was recorded, unless the participant’s scan could be obtained from the database of the institute. The MRI data was recorded with the 3T Siemens Magnetom Skyra MR scanner (Erlangen, Germany).

### MEG analysis

MEG analysis was performed using the Fieldtrip toolbox (Oostenveld et al., 2011), EEGlab (Delorme & Makeig, 2004), and custom MATLAB scripts.

#### Preprocessing

The raw continuous data was downsampled to 300 Hz and epoched relative to the first cue (from −1.5 s to +10 s). A band-stop filter was applied at 50, 100 and 150 Hz to remove line noise and its harmonics. The data was visually inspected to reject trials with artifacts (e.g., muscle artifacts, SQUID sensor jumps). Next, the data was bandpass filtered between 3 and 30 Hz (Butterworth IIR filter) and ICA was computed (i.e., EEGlab implementation of the infomax ICA algorithm of Bell & Sejnowski (1995)). Finally, the IClabel algorithm (Pion-Tonachini et al., 2019) was used to classify components in the categories Brain, Muscle, Eye, Heart, Line Noise, Channel Noise and Other based on their spatial topography.

#### Sensor-level analysis

First, independent components that were classified as muscle, eyes, heart or channel noise by the IClabel algorithm with a probability higher than 80% were discarded. Further analysis was performed at sensor level after back-projecting the remaining components. We computed the planar representation of the MEG field distribution from the single-trial data using the nearest-neighbor method. This transformation facilitates the interpretation of the sensor level data as it makes the signal amplitude maximal above its source (as implemented in Fieldtrip functions *ft_meg_planar* and *ft_combine_planar*). The power spectrum of each channel between 5 and 25 Hz was obtained using a multitaper frequency transformation (*ft_freqanalysis*). This transformation was done separately for the fixation (1 s) and the memory delay period (1 s window centered in the delay period). The data was zero-padded to 5 s to obtain a frequency resolution of 0.2 Hz. Alpha band power was estimated as the mean power values between 8 and 14 Hz across trials. Individual alpha power and frequency were estimated using the MATLAB *findpeaks* algorithm (i.e., maximum peak between 8 and 14 Hz).

#### Component-level analysis

A series of conditions were imposed to select posterior oscillatory components in the alpha range. First, independent components had to be classified as brain components by the IClabel algorithm with a probability higher than 80%. Second, independent components had to show a maximum peak in the alpha range (as detected with the MATLAB *findpeaks* function). Third, independent components needed to project to a single dipole in occipital or parietal cortex. For that purpose, the whole brain was scanned with a single dipole to find the location where the dipole model was best able to explain the topography of each independent component (*ft_dipolefitting*). The source localization of dipoles was done using individual T1-weighted anatomical images of the participants’ brains. For that, individual MRIs were first normalized in MNI space (*ft_volumenormalise*) and segmented (*ft_volumesegment*). Then a realistic single-shell headmodel was computed (*ft_prepare_headmodel)* based on the surface mesh obtained from the segmented MRI (*ft_prepare_mesh*) (Nolte, 2003). Finally, in order to automatically identify dipoles that were located in occipital and parietal cortices we used the AAL atlas (Tzourio-Mazoyer et al., 2002) available in the Fieldtrip toolbox (Oostenveld et al., 2011). The frequency transformation of independent components was identical to the one used for sensor level analysis. The only difference is that in order to estimate alpha power per condition in single trials, the individual alpha band was previously defined per component based on its average power spectrum (across trials; Figure 1B). This approach was adopted to compensate for the lower signal-to-noise ratio of oscillatory activity when estimated in single trials. A similar approach was adopted to estimate peak frequency in single trials. In this case, independent components were first filtered (MATLAB function *firl1*) around their individual alpha band, after which instantaneous frequency was estimated with the method developed in Cohen (2014). In short, instantaneous frequency was computed by multiplying the first temporal derivative of the phase angle time series (extracted through its Hilbert transform) by the sampling rate and dividing it by 2π. Then, a median filter was applied to the instantaneous frequency time series (10 steps between 10 and 400 ms) in order to attenuate non-physiological frequency jumps. Instantaneous frequency was averaged within the fixation period and the memory delay (1-s windows) to get an estimation of peak frequency in each period.

### Statistical analysis

#### Behavioral data

Repeated-measures ANOVA was performed with the JASP software (Love et al., 2019), post-hoc paired samples t-test were performed in MATLAB R2021a. The effect size was estimated as Eta squared (n^2^) for the ANOVA and as Cohen’s d for the t-tests.

#### MEG data

The comparison of MEG parameters of interest between conditions was performed using paired samples and independent samples t-tests (MATLAB R2021a implementation). For the comparison of the parameters of interest in independent components at single-trial level (i.e., power/frequency during memory retentions vs. delay), we employed Wilcoxon signed rank test (MATLAB R2021a) to minimize the effect of possible outliers because they are more likely to occur when analyzing single trials (Cohen & Cavanagh, 2011). In order to assess a possible relationship between an MEG parameter and accuracy, a median split approach was adopted. In short, trials were divided into two groups based on the median of the MEG parameter (i.e., high and low alpha power) and mean accuracy was compared between ‘high’ and ‘low’ trials at group level (paired-samples t-tests). We corrected for multiple comparisons using the False Discovery Rate (FDR) method (Benjamini & Hochberg, 1995). Effect size was estimated through Cohen’s d estimate.

## Results

### Behavioral performance

Mean accuracy was 87.5% across conditions (SD = 19.4). Repeated-measures ANOVA revealed a significant main effect of the factor distractor (F(1,32) = 11.54; p = 0.002; n^2^= 0.102). Post-hoc paired sample t-tests showed that accuracy was greater for the no distractor than for the distractor condition (t(32) = −3.39; p = 0.0018; d = 0.59). No main effect of rule (i.e., stay versus switch) or rule by distractor interaction was found.

Mean reaction time was 698 ms across conditions (SD = 255). Repeated-measures ANOVA revealed a main effect of rule (F(1,32) = 8.79; p = 0.006; n^2^= 0.12). Post-hoc paired sample t-tests showed that reaction time was shorter for stay versus switch rules (t(32) = 2.96; p = 0.0057; d = 0.51). No effect of distractor nor rule by distractor interaction was found.

### Posterior individual alpha increases in power and decreases in frequency during memory retention

No significant differences between conditions in alpha power were found when estimated using an a priori definition of the alpha band (8–14 Hz; Figure 2A, top plot). However, individual alpha peak power showed a significant increase during the memory delay in posterior and right frontocentral sensors (p < 0.05 after FDR correction; Figure 2A, middle plot). In addition, individual alpha frequency decreased significantly in posterior and frontal sensors (p < 0.05 after FDR correction; Figure 2A, bottom plot). Hence, some posterior sensors showed a significant modulation in both individual alpha peak power and frequency (Figure 2B). No significant distractor effect (comparison of distractor vs. no distractor conditions) or relation to accuracy (median split approach) were found in either individual alpha power or frequency at sensor level.

**Figure 2.**
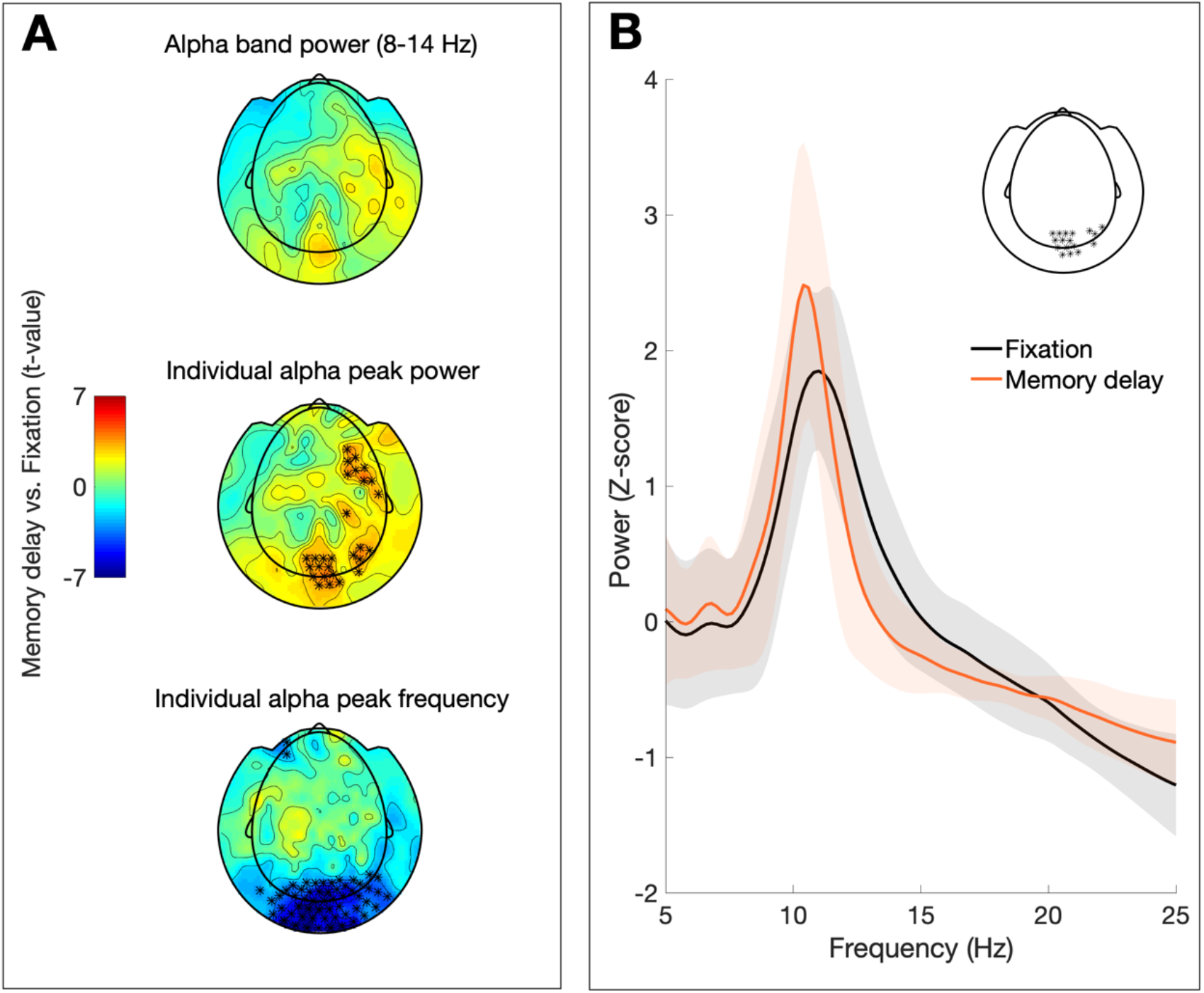
Sensor level analysis. A) Topographic plots depicting the t-values from the condition comparison (memory delay vs. fixation) in alpha band power (top panel), individual alpha peak power (middle) and individual alpha peak frequency (bottom). Significant differences (p < 0.05 after FDR correction) are marked with asterisks. B) Mean power spectrum for fixation (black graph) and memory delay (orange) of sensors showing significant changes in individual alpha peak power and frequency (shaded area depicts standard deviation across subjects; sensors included in spectra indicated in inset).

### ICA reveals two distinct alpha components based on their power modulations during the memory delay

In order to assess whether the reported changes in posterior alpha power reflect the activity of one or several brain rhythms, we performed the same condition comparisons that we performed at sensor level using independent posterior alpha components (Figure 1B). We found a total of 170 posterior alpha components based on their topography at sensor level, their spectrum, and their estimated source. The power of 111 components was significantly modulated during the memory delay relative to fixation (p < 0.05 after FDR correction; mean number of components per subject = 3.3; SD = 2.9). Unlike sensor-level analysis, the comparison of posterior alpha components in single subjects revealed both increases and decreases in alpha power during the memory delay. Specifically, the power of 68 alpha components showed a significant increase during memory delay relative to fixation (Alpha1), while the power of 43 alpha components showed a significant decrease (Alpha2; Figure 3A).

**Figure 3.**
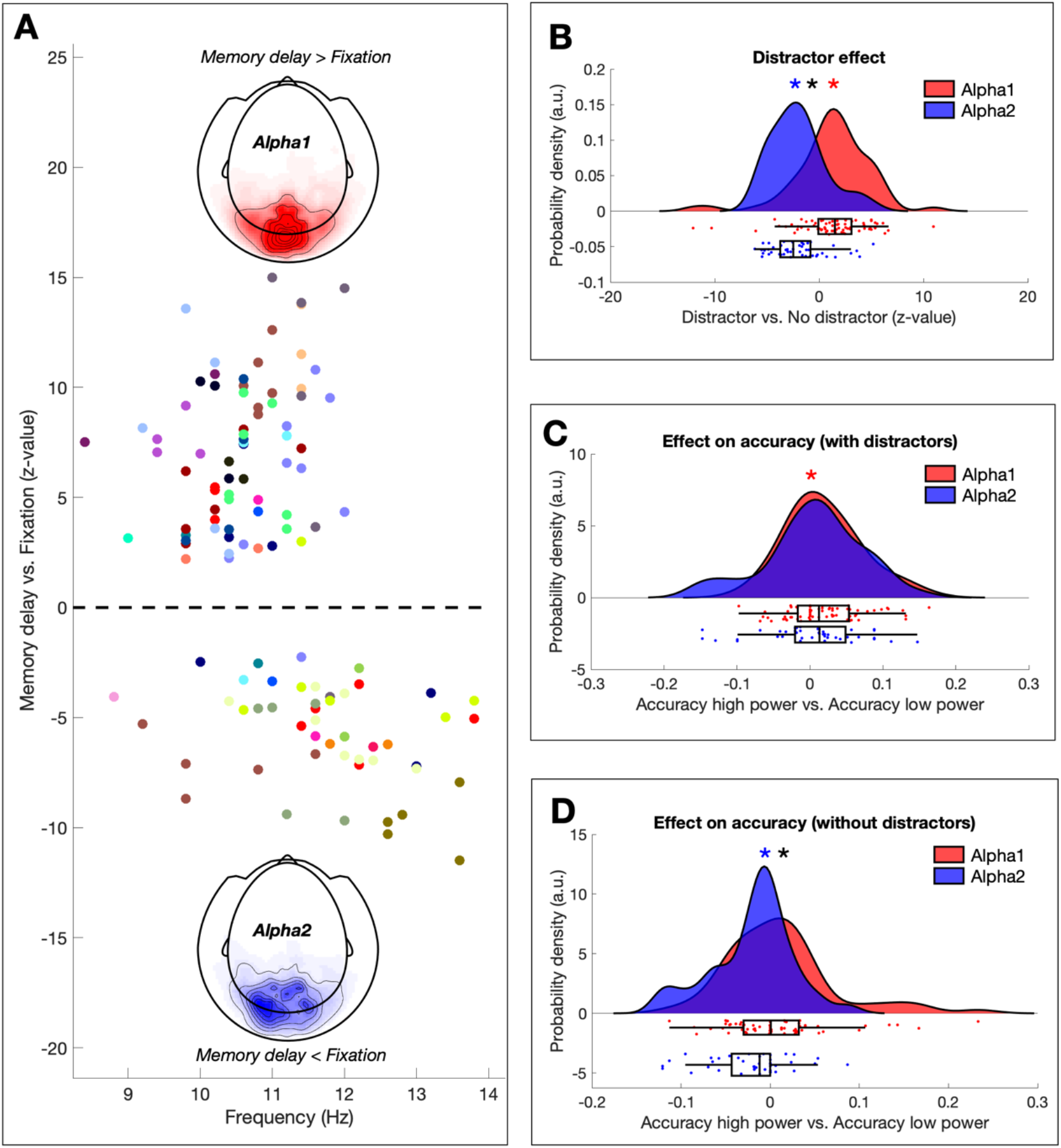
Power modulations of independent alpha components. A) Power changes during the memory delay relative to fixation. Z-values (representing the memory retention effect in each component) are plotted as a function of their peak frequency (each color codes for a different subject). Components showing a significant increase in power during the memory delay were denominated Alpha1 while components showing a significant decrease were denominated Alpha2. The mean topography of the power change is plotted separately for Alpha1 (red) and Alpha2 (blue) components. B) Differential distractor-related power modulations of Alpha1 and Alpha2 components. Z-values represent the distractor effect in individual components. At group level, Alpha1 components (red) showed significantly more power in the presence of distractors while Alpha2 components (blue) showed significantly less. Colored asterisks mark statistical significance (p < 0.05) of the Alpha1 (red) or Alpha2 (blue) distributions against 0 (one-sample t-test). Black asterisks mark significant differences (p < 0.05) between Alpha1 and Alpha2 distributions. C) Effect of power modulations on accuracy in the distractor condition. Accuracy was compared for high power and low power trials (median split), showing a significantly greater accuracy for trials with high Alpha1 power in the distractor condition. D) Same as C for the no-distractor condition, showing a significantly lower accuracy for trials with high Alpha2 power in the no-distractor condition.

In addition, we assessed whether the frequencies of Alpha1 and Alpha2 components were differentially modulated during memory delay. However, we did not find significant differences between Alpha1 and Alpha2 components in frequency modulations associated with memory retention (t(109) = −1.45; p = 0.14; d = 0.28). Both Alpha1 and Alpha2 components significantly decreased in frequency during the memory delay relative to fixation (t(67) = −8.18, p < 0.001, d = 1.00; t(42) = −6.98, p < 0.001, d = 1.07).

In summary, single-subject analysis of posterior alpha components demonstrates the existence of at least two distinct rhythms (Alpha1 and Alpha2) based on their opposite power modulations during memory retention.

### Alpha1 and Alpha2 show opposite distractor-related power modulations

In order to assess whether the power of Alpha1 and Alpha2 components was differentially modulated in the presence of distractors, we first estimated the distractor effect in individual components (distractor vs. no distractor) through Wilcoxon signed rank tests. Then we tested at group level whether the z-values of Alpha1 and Alpha2 components differed significantly from 0 (one sample t-test) and from each other (independent samples t-test). We found that alpha components that increased in power during the memory delay (i.e., Alpha1) showed a significant power increase in the presence of distractors (t(67) = 2.66; p = 0.0097; d = 0.32) while alpha components that decreased in power during the memory delay (i.e., Alpha2) showed a significant power decrease in the presence of distractors (t(42) = −5.78; p < 0.001; d = 0.88). Hence, Alpha1 and Alpha2 showed opposite and significantly different (t(109) = 5.42; p < 0.001; d = 1.05) distractor-related power modulations (Figure 3B).

### Alpha1 and Alpha2 power modulations have an opposite relation to accuracy

In order to assess the behavioral relevance of Alpha1 and Alpha2 power modulations, we compared the accuracy between trials with high and low alpha power during the delay (% change from fixation). Since the presence of distractors was associated with lower accuracy, we performed this analysis for distractor and no-distractor conditions separately. For the distractor condition, accuracy was significantly higher for trials showing greater Alpha1 power (t(67) = 2.97; p = 0.0041; d = 0.36), while no differences were found in Alpha2 power (t(42) = 0.54; p = 0.58; d = 0.08). However, power-related differences in accuracy between Alpha1 and Alpha2 did not reach statistical significance (t(109) = 1.18; p = 0.24; d = 0.23). For the no-distractor condition, accuracy was significantly higher for trials with lower Alpha2 power (t(42) = −2.79; p = 0.0078; d = 0.42), while no significant difference was found in Alpha1 power (t(67) = 1.42; p = 0.15; d = 0.17). In this case, power-related differences in accuracy between Alpha1 and Alpha2 components did reach statistical significance (t(109) = 2.76; p = 0.0067; d = 0.53).

In summary, in the presence of visual distractors, better accuracy was associated with higher Alpha1 power, while in the absence of visual distractors, better accuracy was associated with lower Alpha2 power.

### Alpha1 and Alpha2 components differ in their spatiospectral characteristics

In order to assess whether Alpha1 and Alpha2 rhythms differ in their spatiospectral characteristics, we compared three different spectral parameters (peak frequency, peak width, and relative amplitude; Figure 4A) and the location of their estimated main source through dipole fitting (x, y and z axes; Figure 4B). This analysis revealed that Alpha1 components tended to have a lower peak frequency (t(109)= −6.00; p < 0.001; d = 1.16), a narrower peak width (t(109)= −2.43; p = 0.016; d = 0.47), a greater relative peak amplitude (t(109)= 5.68; p < 0.001; d = 1.10), and a more central source estimation (t(109)= 2.86; p = 0.005; d = 0.55) than Alpha2 components (Figure 4C). No significant differences were found between the source estimation of Alpha1 and Alpha2 components in the other two axes (i.e., ventral to dorsal and posterior to anterior).

**Figure 4.**
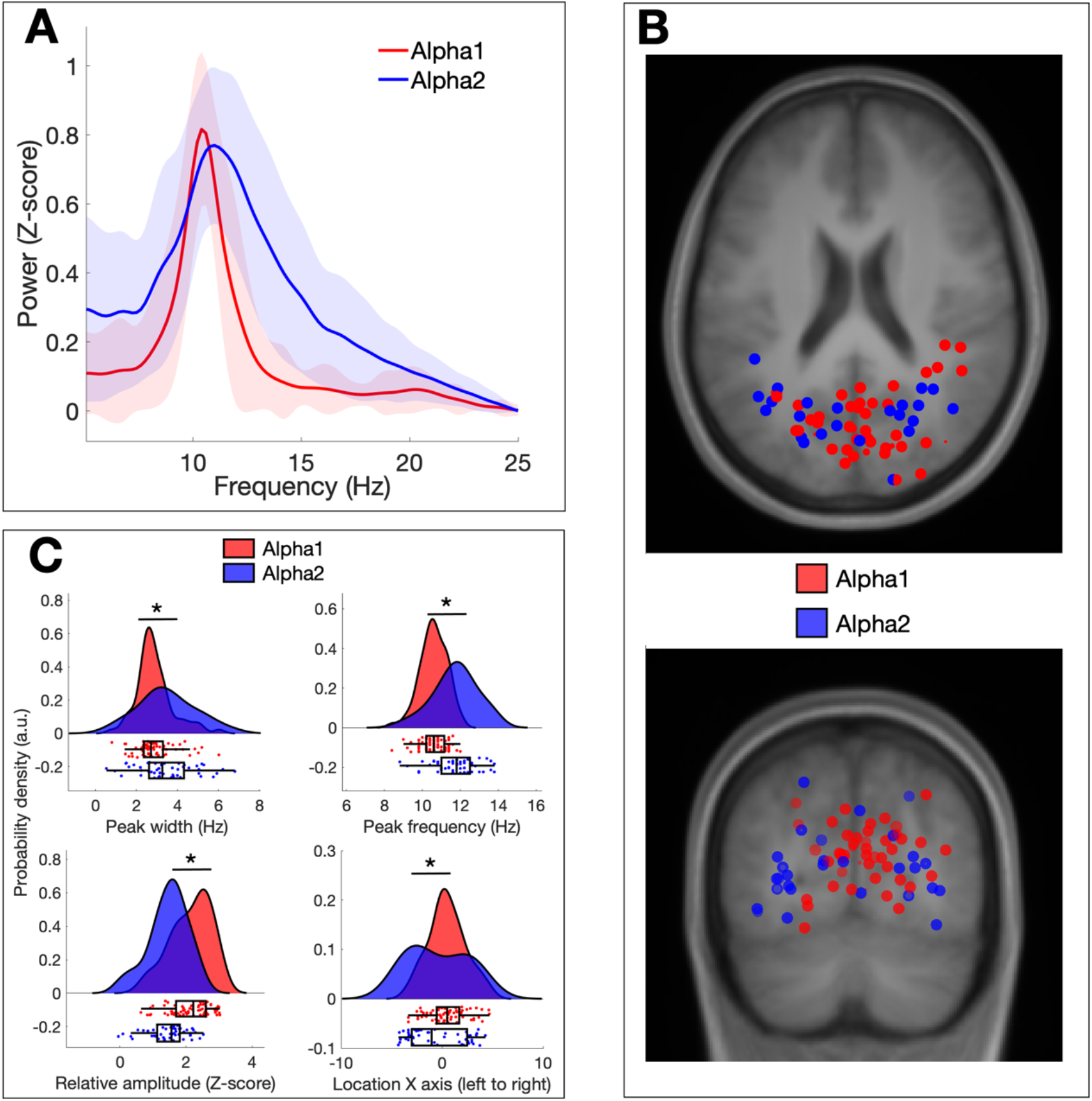
Spatiospectral differences between Alpha1 and Alpha2 components. A) Average spectrum of Alpha1 and Alpha2 components across the complete trial (shaded area depicts standard deviation). B) Source localization of Alpha1 (red) and Alpha2 (blue) components as estimated through dipole fitting. The top panel shows the horizontal plane while the bottom panel shows the coronal plane. C) Plots depicting significant differences in four spatiospectral parameters: peak width (top left panel), peak frequency (top right), relative amplitude (bottom left) and location (bottom right).

## Discussion

In this study we used ICA to examine whether posterior alpha power modulations during visual working memory reflect the dynamics of one or several brain rhythms. We recorded MEG while participants (N = 33) performed a visual working-memory task in which one out of four spatial directions had to be remembered for a short period of time. Task difficulty was modulated by introducing visual distractors during memory retention. Group analysis at sensor level suggested that posterior alpha consists of a single oscillator that increases in power and decreases in frequency during memory retention. In contrast, the analysis of independent components in single subjects revealed the existence of an alpha rhythm that increases in power during the memory delay (Alpha1), and an alpha rhythm that decreases in power during the memory delay (Alpha2). Interestingly, the power of Alpha1 and Alpha2 rhythms was differentially modulated by the presence of distractors (Alpha1 increased in power while Alpha2 decreased), and had an opposite relationship with accuracy (positive for Alpha1 and negative for Alpha2). In addition, Alpha1 and Alpha2 rhythms differed significantly in their spatiospectral characteristics. Specifically, Alpha1 rhythms showed a lower peak frequency, a narrower peak width, a greater relative peak amplitude and a more central source than Alpha2 rhythms. Thus, our results show that modulations in posterior alpha oscillations during memory retention reflect the dynamics of at least two distinct brain rhythms with different functions and spatiospectral characteristics.

Previous literature is highly inconsistent regarding the direction of alpha power modulations during visual memory retention. A recent systematic review has shown that from 56 M/EEG studies, 30 report a significant alpha power increase during memory retention, 21 report a significant decrease and 5 show either mixed or null results (Pavlov & Kotchoubey, 2020). Based on our results, we speculate that the lack of consistency in previous literature could be traced to the commonly adopted analytical approach of computing alpha power by averaging over a predefined frequency band (e.g., 8–14 Hz). It has been shown that alpha power modulations can be easily confounded by frequency changes if peak detection is not performed (Donoghue et al., 2021). This is important because alpha peak frequency not only varies considerably between subjects (Doppelmayr et al., 1998), but also within subjects in a task-dependent manner (Haegens et al., 2014; Rodriguez-Larios & Alaerts, 2019; Samaha & Postle, 2015). In line with this, our results showed that the increase in alpha peak power at sensor level during the memory delay was accompanied by a frequency decrease (Figure 2). Given this frequency shift, significant alpha power modulations could have been reported in different directions depending on the a priori definition of the alpha band (e.g., 8–12 vs. 9–13 Hz). Consequently, we hypothesize that performing peak frequency detection could solve at least some previous inconsistencies regarding the direction of alpha power modulations during visual memory retention (Pavlov & Kotchoubey, 2020).

Although peak detection at sensor level allows disentangling alpha power and frequency modulations, it cannot determine whether the reported changes reflect the activity of one or multiple brain rhythms (Schaworonkow & Nikulin, 2019). In line with previous studies (Barzegaran et al., 2017; Benwell et al., 2019; Gulbinaite et al., 2017; Sokoliuk et al., 2019), we demonstrate the existence of two different alpha rhythms in posterior cortex. Specifically, the analysis of independent components in single subjects revealed that a faster alpha rhythm (Alpha2) decreased in power during memory retention while a slower alpha rhythm (Alpha1) increased in power. In this regard, it is important to underline that differential power modulation of two alpha rhythms with different peak frequencies could lead to apparent frequency modulations at sensor level (Donoghue et al., 2021). Therefore, we cannot rule out the possibility that previously reported frequency changes in posterior alpha during different cognitive tasks (Angelakis et al., 2004; Babu Henry Samuel et al., 2018; Haegens et al., 2014; Rodriguez-Larios & Alaerts, 2019) are actually reflecting power changes of two (or more) alpha rhythms with different spatiospectral characteristics.

The existence of alpha rhythms that increase and decrease in power during memory retention depending on their spatial origin is in line with prevailing theories of alpha function (Jensen & Mazaheri, 2010; Klimesch et al., 2007). According to these theories, alpha power reflects local inhibition and therefore should increase in task-irrelevant areas and decrease in task-relevant areas. In the context of memory retention, it can be predicted that brain regions that are relevant to the transient maintenance of visual information would show alpha decrease (disinhibition). On the other hand, brain regions that are irrelevant for memory maintenance (and could potentially interfere with the task) would show alpha increase (inhibition). Although source estimation through dipole fitting in M/EEG has to be interpreted with caution (Leahy et al., 1998), the spatial distribution of the two observed types of alpha components suggests that, at least in occipital cortex, Alpha1 rhythms mostly originate in early visual areas while Alpha2 rhythms localize to higher-order areas (Figure 4B). This is supported by their spectral profiles, since higher-order areas are thought to show a more pronounced 1/f trend (Ibarra Chaoul & Siegel, 2021) and higher peak frequency (Lundqvist et al., 2020), i.e., in line with what we see in Alpha2 components when compared to Alpha1 (Figure 4A). Based on these results and previous evidence (De Vries et al., 2018; Popov et al., 2017; Tuladhar, Ter Huurne, et al., 2007), we speculate that Alpha1 power increases reflect the inhibition of lower-order areas involved in visual processing, whilst Alpha2 power decreases reflect the disinhibition of higher-order areas supporting the transient storage of visual information.

If Alpha1 and Alpha2 rhythms during memory retention reflect the inhibition and disinhibition of task-irrelevant and task-relevant areas respectively, we would expect that behavioral performance improves when Alpha1 power increases and Alpha2 power decreases. Interestingly, Alpha1 power increases and Alpha2 power decreases were associated with better accuracy in different experimental conditions. Specifically, Alpha1 power was positively associated with accuracy only in the presence of distractors, while Alpha2 power was negatively associated with accuracy only in the absence of distractors. We hypothesize that behavioral performance in distractor and no-distractor conditions depends on different factors. On the one hand, incorrect responses in the distractor condition might mostly be due to the interference of visual distractors. In this scenario, Alpha1 power increases become predictive of behavior because it inhibits areas involved in visual processing in order to avoid interference during memory retention. On the other hand, in the condition without distractors, incorrect responses might predominantly be caused by lapses of attention due to mind wandering and/or drowsiness (Andrillon et al., 2019, 2021; Braboszcz & Delorme, 2011; Rodriguez-Larios & Alaerts, 2020). Previous literature suggests that lapses of attention involve reduced cortical processing of external events (Smallwood et al., 2008). If an external event is not properly processed by the brain during an attentional lapse, its content cannot be maintained in working memory. Hence, we can expect that the recruitment of cortical areas supporting short-term memory retention (i.e., Alpha2 desynchronization) is less pronounced (or even absent) during an attentional lapse because there is little or no information to be retained. Nonetheless, it is important to note that the reported differences in accuracy depending on Alpha1/Alpha2 power modulations must be interpreted with caution due to the small number of incorrect trials here (mean accuracy was 87% across conditions). In order to overcome this limitation in future work, it would be important to assess the here reported effects with a more difficult task or by adjusting task-difficulty at an inter-individual level.

In line with previous literature, our results show that ICA is a powerful analytical tool that can be efficiently used to isolate brain rhythms of interest (Benwell et al., 2019; Debener et al., 2005; Wagner et al., 2018). Unlike other source localization techniques, ICA does not require a priori definition of the specific spatial location, and ensures that the analyzed time series are statistically independent (thereby minimizing the possibility that they reflect the mix of two or more rhythms; Delorme et al., 2012). The spatial separation of different posterior alpha rhythms through ICA in single subjects could resolve some of the previous inconsistencies in the literature concerning the role of alpha phase, power and frequency in cognition (Michail et al., 2021; Pavlov & Kotchoubey, 2020; Samaha et al., 2020; Zazio et al., 2021). Similarly, separating different posterior alpha rhythms could allow us to understand why some neurofeedback and neurostimulation protocols do not have the expected effect in some subjects (Orendáčová & Kvašňák, 2021). If we tune the neurofeedback/stimulation parameters (and/or assess their effects) at sensor level, we cannot know whether we are modulating the power or frequency of one or several alpha rhythms. As different alpha rhythms could be more prominent in different subjects due to inter-individual differences in brain anatomy and functional specialization, this might have a key impact on the effects of their modulation (Duffau, 2017; Mikkonen et al., 2020).

In conclusion, our results show that posterior alpha dynamics during memory retention reflect the activity of at least two brain rhythms with distinct functions and spatiospectral characteristics. Alpha1 rhythms increased in power during memory retention and in the presence of visual distractors, while Alpha2 rhythms showed the opposite power modulations. In addition, Alpha1 and Alpha2 rhythms had an opposite relationship with accuracy (positive for Alpha1 and negative for Alpha2). Lastly, Alpha1 and Alpha2 differed significantly in several spectral parameters (peak frequency, peak width and relative amplitude) and in the location of their estimated main source. In the light of previous results and theoretical accounts (Haegens et al., 2021; Iemi et al., 2022; Jensen & Mazaheri, 2010; Klimesch et al., 2007), we hypothesize that during memory retention, Alpha1 rhythms increase in power to inhibit visual processing while Alpha2 rhythms decrease in power to disinhibit areas supporting the short-term storage of visual information.

## Notes

**Funding Sources** This work was supported by NWO Vidi grant 016.Vidi.185.137 and by NIH grant R01-MH123679.

**Conflict of interest statement** The authors declare no conflict of interest.

### Competing Interest Statement

The authors have declared no competing interest.

